# SH3KBP1/CIN85, a new actor of ER-phagy in muscle

**DOI:** 10.64898/2026.07.15.737746

**Authors:** Marine Daura, Elina Vergara, Leslie Andromaque, Agathe Leddet, Emilie Christin, Céline Malleval, Vincent Gache, Carole Kretz-Remy

**Affiliations:** Université Lyon1, CNRS, INSERM, INMG-PGNM, UMR5261, U1315, Lyon, France

**Keywords:** Autophagy, CIN85, CKAP4, Climp63, endoplasmic reticulum, reticulophagy, Ruk, striated skeletal muscle

## Abstract

The endoplasmic reticulum (ER) and its muscle-specialized form, the sarcoplasmic reticulum (SR), are crucial organelles in muscle cells, involved notably in protein synthesis, calcium regulation and muscle contraction. A well-known process involved in ER remodeling and homeostasis is ER-phagy, also called reticulophagy, a selective form of autophagic process in which ER-phagy receptors mediate the delivery of ER portions to lysosomes for degradation. SH3KBP1 is an adaptor protein involved in membrane trafficking. Recently, it was shown to control ER morphology and SR formation in striated skeletal muscle. In this study, we demonstrate that SH3KBP1 can bind to LC3B and CKAP4 proteins, bridging ER to autophagosome membranes, and is degraded by autophagy, in developing muscle fibers. Moreover, SH3KBP1 down-regulation impacts basal autophagy efficiency and ER-phagy stimulation; it also impairs the turnover of numerous ER-resident proteins. Our work highlights a new role for SH3KBP1 as a soluble ER-phagy receptor in striated skeletal muscle.

## Introduction

Macroautophagy (hereafter autophagy) is a catabolic process during which double membrane organelles called autophagosomes (APs) are formed, engulfing parts of cytoplasm; they next fuse with acidic lysosomes to form autolysosomes (ALs) whose content is degraded by lysosomal hydrolases and recycled into the cytoplasm [1,2]. This mechanism, often regulated by mTOR (mechanistic target of rapamycin kinase) and/or AMPK (5’ adenosine monophosphate-activated protein kinase), relies on the sequential recruitment of ULK1 (unc51 like autophagy activating kinase 1) and PI3KC3 (class III phosphatidyl-inositol 3-kinase)-BECN1/beclin1 complexes, ATG9 (autophagy related 9) vesicles, WIPIs (WD repeat domain phosphoinositide interacting proteins) and ATG12—ATG5 conjugation systems, allowing the lipidation of ATG8 (MAP1LC3/LC3 [microtubule associated protein light 1 chain 3] and GABARAP (GABA type A receptor-associated protein) protein families at the site of phagophore formation [3]. Autophagy constitutively occurs at basal level to maintain the turnover of the cytoplasmic components. Under stress conditions, such as nutrient deprivation, bulk (non-selective) autophagic activity is induced as a recycling process, generating metabolites and sustaining energy production [4]. Autophagy can also specifically target protein aggregates, lipid droplets, invading pathogens or damaged organelles such as mitochondria or endoplasmic reticulum (ER) [5,6]. Selective autophagy receptors ensure this specificity through their ability to bind both cargoes and members of phagophore-associated ATG8 or GABARAP protein families [7]. The use of a great variety of autophagy receptors and a conserved core autophagic machinery enable cells to adapt, restore their homeostasis and optimize their energy expenditure.

The ER plays a central role in autophagy as a major membrane source for autophagy initiation and APs biogenesis [8]. It is a highly dynamic organelle involved in protein quality control, carbohydrate metabolism, calcium signaling or lipid biosynthesis [9]. The ER consists of a complex network of membrane tubules and flattened cisternae (called sheets) that are constantly remodeled in response to cellular demands such as during mitosis, cell differentiation, the unfolded protein response (UPR) or exposure to chemical stresses [10]. Its size and activity are regulated by ER-phagy (also called reticulophagy), a selective autophagic process that degrades ER fragments and content, thereby contributing to ER reshaping. Consistently, inhibition of autophagy leads to ER expansion and dilatation [8,11]. In mammals, ER-phagy process is mediated by ER-phagy receptors that contain LC3-interacting region (LIR) motifs, and act as adaptors linking ER to APs membranes. As a consequence, ER-phagy receptors are themselves sequestered into APs and thus degraded by autophagy. To date, twelve ER-phagy receptors have been described, which target specific ER subdomains and coordinate stress responses in diverse cell types or tissues [8]. These receptors are classified into two categories: i) membrane-bound ER-phagy receptors, which are anchored in the ER membrane through transmembrane domains and may contain reticulon-homology domains (RHD) and ii) cytosolic ER-phagy receptors, which are recruited to the ER through interactions with ER resident transmembrane proteins [8,12]. Only portions of the ER are selectively targeted for fragmentation and delivery to lysosomes, although the molecular mechanism underlying this process remains not yet fully understood. Post-translational modifications, receptor clustering or oligomerization appear to be key steps in this process [8,13,14]. Moreover, it is very likely that a cooperation with ER-resident adaptors and molecular chaperones is needed since most ER-phagy receptors lack luminal domains. Together, they facilitate cargo segregation, membrane shaping and shedding. ER-phagy can be activated by pleiotropic signals such as cellular stresses or chemical compounds that induce a global cellular response, including diverse pathways and also non-selective autophagy. ER-phagy can also be specifically induced, without affecting other cytosolic components, by «ER-centric activators such as accumulation of misfolded polypeptides, ER stresses or infections [8].

ER remodeling is essential in most cell types but is particularly prominent in skeletal muscle, where part of the ER differentiates into the specialized sarcoplasmic reticulum (SR), a complex tubular network essential for transducing electrical impulses into contractile force through the release and recapture of stored calcium ions. During myogenesis, extensive remodeling is required to organize ER subdomains, a process in which ER-phagy plays an important role. As myocytes fuse to form multinucleated myofibers, heterotypic ER membranes fusion occurs concomitantly with increased ER-phagy activity [15,16].

SH3KBP1/CIN85 (SH3 domain containing kinase binding protein 1/Cbl-interacting protein of 85kDa) is a ubiquitously expressed adaptor protein belonging to the CMS family (Cas Ligand with multiple SH3 domains) [17]. Its N-terminal (N-ter) region contains three SH3 domains followed by a proline-rich domain, enabling interactions with numerous partners and assembly of diverse multiprotein complexes [18,19]. Moreover, a coiled-coil C-terminal (C-ter) part allows both tetramerization and interaction with membrane lipids [20]. SH3KBP1 is localized in the cytoplasm, mainly associated to vesicular structures including endocytic vesicles, the ER-Golgi intermediate compartment (ERGIC), the cis-Golgi network and COP-I-coated vesicles [21]. Due to its adaptor function, SH3KBP1 is involved in signal transduction, endocytic trafficking, degradation and rearrangement of cytoskeleton structures [17,22]. We recently demonstrated that SH3KBP1 is a regulator of myofiber homeostasis, controlling myoblast fusion, myotubes elongation and nuclear positioning, as well as ER and SR integrity in mature myofibers. In addition, we observed that intramuscular downregulation of SH3KBP1 in mice induces muscle atrophy [23], which raises the question of which pathways are regulated by this protein.

Here, we show that SH3KBP1 regulates basal autophagic and ER-phagic activities. We also demonstrate that SH3KBP1 is degraded by autophagy, contains LIR motifs and interacts with both LC3B and the ER-transmembrane proteins CKAP4/CLIMP63 (cytoskeleton associated protein 4/63kDa cytoskeleton-linking membrane protein), pointing a direct role of SH3KBP1 in ER cargo recognition. These interactions are modulated upon autophagy and ER-phagy induction. We thus propose that SH3KBP1 is a novel soluble ER-phagy receptor in striated skeletal muscle cells, our findings expanding the repertoire of described mechanisms that govern selective ER turnover.

## Results

### SH3KBP1 supports basal autophagic flux

Downregulation of SH3KBP1 adaptor protein was shown to induce a disorganization of the perinuclear ER in developing myofiber [23]. Since autophagy is a major process involved in the maintenance and remodeling of ER architecture and given that ER is important for membrane supply and AP formation [8], we first sought if SH3KBP1 was involved in the regulation of basal autophagy in muscle cells.

To address this question, the double-tandem fluorescent reporter mammalian red fluorescent protein-green fluorescent protein-LC3 (mRFP-GFP-LC3) [24] was transfected in C2C12 myoblasts stably expressing a control shRNA (shCTRL) or an shRNA targeting SH3KBP1 (shSH3KBP1) [23]. The cells were then treated or not with the commonly used autophagic flux inhibitor Bafilomycin A1 (BafA1) [25]. Next, autophagic flux was quantified as the ratio of GFP^+^RFP^+^ vesicles to total-RFP^+^ vesicles (Figure 1A-B). In untreated shCTRL cells, RFP^+^-only vesicles (ALs) were more abundant than GFP^+^RFP^+^ vesicles (APs), consistent with an active basal autophagic flux under steady-state conditions. Upon BafA1 treatment, the number of GFP^+^RFP^+^ vesicles markedly increased compared to the dimethyl sulfoxide (DMSO, vehicle) condition, while RFP^+^-only vesicles were almost completely absent, indicating, as expected, a blocked APs maturation (Figure 1A). Accordingly, quantification revealed an increase in the GFP^+^RFP^+^: total-RFP^+^ ratio (1.8-fold change) (Figure 1B, light grey). In contrast, at steady state, untreated shSH3KBP1 myoblasts, in which SH3KBP1 is downregulated (Figure S1A), displayed a higher number of GFP^+^RFP^+^ vesicles and a reduced number of RFP^+^-only vesicles compared to untreated shCTRL cells (Figure 1A). Accordingly, the GFP^+^RFP^+^: total-RFP^+^ ratio was significantly increased in non-treated (DMSO) shSH3KBP1 cells compared to shCTRL cells (Figure 1B). Upon BafA1 treatment, GFP^+^RFP^+^ vesicles still accumulated in shSH3KBP1 cells, indicating that APs can still be generated but to a smaller extent (1.3-fold change) (figure 1 A-B, dark grey). Altogether, these data indicate that basal autophagic flux is altered in C2C12 cells under-expressing SH3KBP1; indeed, the increased APs levels at steady state in shSH3KBP1 cells and the reduced accumulation of APs upon lysosomal inhibition in these cells suggests an impaired AP maturation/progression along the flux.

**Figure 1.**
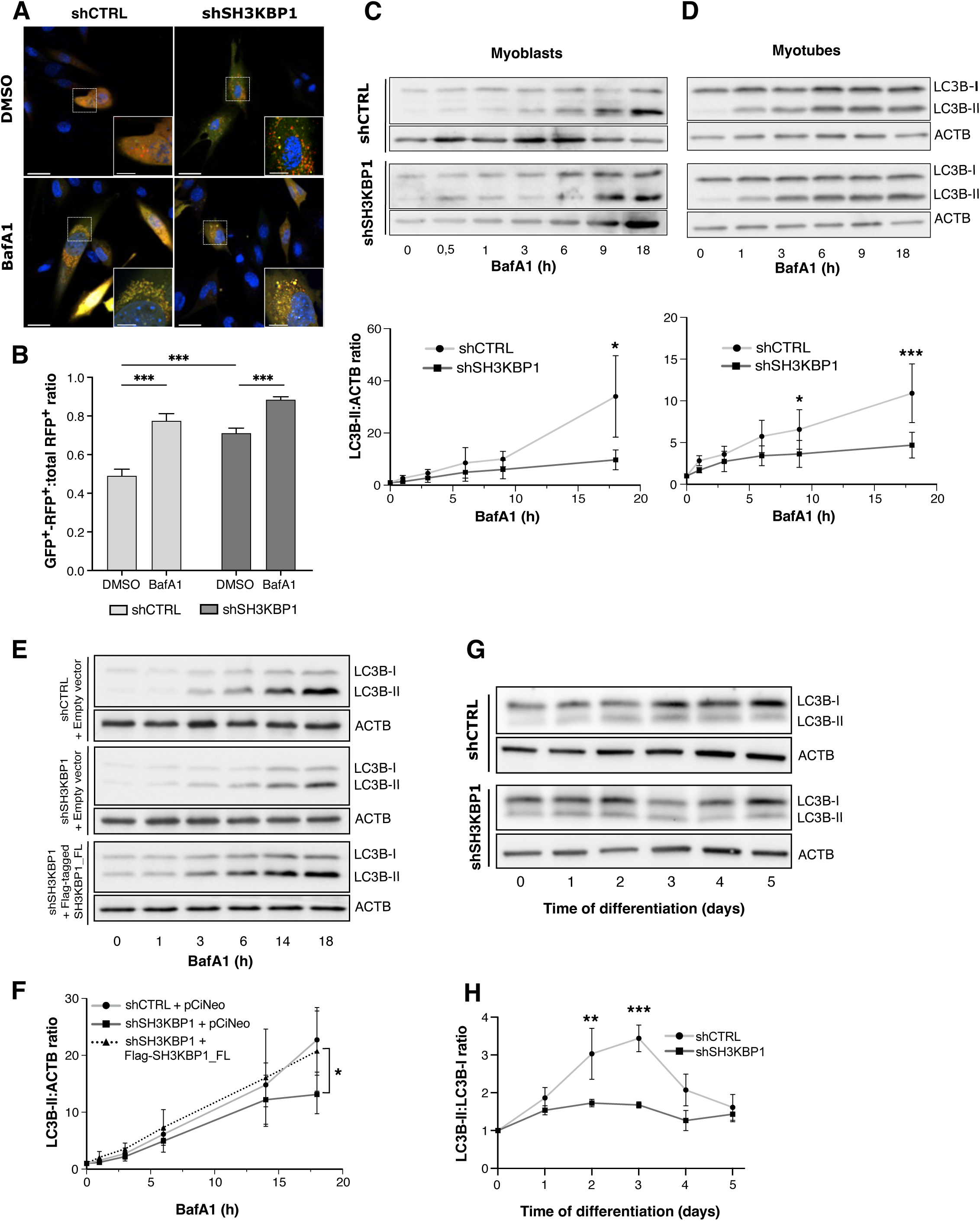
SH3KBP1 downregulation affects basal autophagy in muscle cells. (**A**) Representative confocal images of C2C12 myoblasts stably expressing control shRNA (shCTRL) or a SH3KBP1-targeting shRNA (shSH3KBP1) and transfected with the tandem fluorescent reporter mRFP-GFP-LC3. The cells were treated with DMSO (vehicle) or BafA1. GFP^+^RFP^+^ vesicles (yellow) correspond to APs, whereas RFP^+^-only vesicles (red) correspond to ALs. Scale bar: 30 µm (overview images); 10μm (insets). **B**) Quantification of autophagic activity expressed as the ratio of GFP^+^RFP^+^ vesicles on total RFP^+^ vesicles in shCTRL and shSH3KBP1 C2C12 cells; n=3 independent experiments. (**C and D**) Representative western blot analysis showing the kinetics of LC3B-II accumulation in shCTRL and shSH3KBP1 C2C12 myoblasts (**C**) or in 5-days differentiated myotubes (**D**) treated with 100 nM BafA1 for the indicated periods of time and the corresponding quantification of LC3B-II levels normalized to ACTB levels; n= 3 independent experiments. (**E and F**) Rescue experiment. Representative western blot analysis of the kinetics of LC3B-II accumulation in shCTRL and shSH3KBP1 C2C12 myoblasts transfected with an empty vector (pCVM-GFP) or shSH3KBP1 myoblasts re-expressing a full-length flag-tagged SH3KBP1 (Flag-SH3KBP1_FL) and then treated with 100 nM BafA1 for the indicated times (**E**) and its quantification as a ratio between LC3B-II levels and ACTB levels (**F**); n=3 independent experiments. Data are represented as mean +/-SD. Statistical significances between shCTRL and shSH3KBP1 C2C12 cell lines were assessed using Mann-Whitney tests.

To confirm these observations, we measured the autophagic flux by analyzing the kinetics of LC3-II accumulation by western blot in shCTRL and shSH3KBP1 C2C12 cells treated with BafA1 from 0 to 18 h [26]. LC3-II progressive accumulation over time reflects APs formation and autophagic flux. In C2C12 myoblasts, LC3B-II levels increased similarly in shCTRL and shSH3KBP1 cells lines, during the first 9 h of BafA1 treatment. However, beyond this time point, LC3B-II level continued to increase, up to 18 h, in shCTRL cells, whereas it practically reached a plateau in shSH3KBP1 cells (Figure 1C, western blot). Quantitative analysis revealed a significant difference between the two cell lines after 18 h of treatment with BafA1, indicating reduced dynamic of LC3B-II accumulation when SH3KBP1 is down-regulated (Figure 1C, graph). Based on these kinetics, we calculated the slope of both curves, as an indicator of the speed of APs formation. This speed was reduced by more than two-fold in shSH3KBP1 compared to shCTRL cells (1.9 in shCTRL versus 0.7 in shSH3KBP1), indicating that APs formation is less efficient in SH3KBP1-underexpressing cells. The same analysis was performed in 5 days-differentiated C2C12 myotubes (Figure S1B). In both shCTRL and shSH3KBP1 myotubes, LC3B-II levels progressively increased during BafA1 treatment (Figure 1D, western blot). However, LC3B-II accumulation tended to reach a plateau after 6 h of treatment in shSH3KBP1 myotubes, whereas it continued to increase up to 18 h of treatment in shCTRL myotubes, further supporting the existence of a reduced autophagic flux in SH3KBP1-underexpressing cells (Figure 1D, graph). Consistently, the speed of APs formation was decreased by approximately 2.5-fold in shSH3KBP1-myotubes compared to control ones (0.7 versus 1.8, respectively).

Finally, we observed that the re-expression of SH3KBP1 in shSH3KBP1 cells, by transfection of a Flag-tagged SH3KBP1_FL construct (Figure S1A), restored the kinetics of LC3B-II accumulation (figure 1E, bottom panel) that normalized to shCTRL one (upper panel). In contrast, transfection of shSH3KBP1 cells with an empty vector (figure 1E, middle panel) did not induce any restoration of LC3B-II accumulation during BafA1 treatment. Quantification of LC3B-II:ACTB (actin) ratios confirmed that LC3B-II accumulation is similar in shCTRL cells and shSH3KBP1 cells expressing Flag-SH3KBP1_FL, whereas it remained markedly reduced in shSH3KBP1 cells (Figure 1F). Hence, these results demonstrate that the observed defects in basal autophagic flux are the specific consequence of SH3KBP1 downregulation. Altogether, these complementary approaches demonstrate that SH3KBP1 is required for efficient basal autophagic flux and proper APs formation and maturation in muscle cells.

Basal autophagy corresponds to the constitutive activity of the autophagic machinery and contributes to cellular quality control through elimination of damaged or superfluous components [27]. Yet autophagy can be strongly induced in response to metabolic or pharmacological stimuli, a process commonly referred to as “bulk” autophagy. To determine whether SH3KBP1 downregulation also impacts bulk autophagy, shCTRL and shSH3KBP1 cells were treated with the autophagy inducer rapamycin (Rapa), the autophagy maturation inhibitor BafA1, or a combination of both drugs during 4 h. As expected, Rapa treatment increased LC3B-II levels in both shCTRL and shSH3KBP1 myoblasts, indicating efficient induction of autophagy upon pharmacological mTORC1 inhibition (Figure S1C). Likewise, BafA1 treatment led to a pronounced accumulation of LC3B-II, reflecting inhibition of APs degradation. Combined Rapa and BafA1 treatments further increased LC3B-II accumulation in shCTRL and shSH3KBP1 cell lines, consistent with the additive effects of autophagy induction and lysosomal blockade (Figure S1C). Quantification of LC3B-II:LC3B-I ratios, normalized to untreated conditions within each cell line, revealed comparable responses to Rapa, BafA1 or combined treatments, in both cell lines (Figure S1D), indicating that SH3KBP1 downregulation does not impair the induction of bulk autophagy nor the ability of cells to accumulate APs under conditions of autophagic stimulation. Altogether, these results show that SH3KBP1 is specifically required for efficient basal autophagic flux in both undifferentiated myoblasts and differentiated myotubes, whereas it is dispensable for the induction of bulk autophagy in response to pharmacological stimuli. Moreover, combined fluorescence-based and biochemical approaches further suggest that SH3KBP1 participates in both AP formation and downstream steps of the autophagic process.

Interestingly, autophagy plays an important role during differentiation processes [28], notably in skeletal muscle differentiation [29]. As we previously observed a transient increase of SH3KBP1 expression during differentiation from myoblasts to myotubes, and given that SH3KBP1 downregulation enhanced myoblasts fusion but not myogenic commitment, [23], we examined autophagic activity during the differentiation of shCTRL and shSH3KBP1 C2C12 cells. In shCTRL cells, the LC3B-II:ACTB ratio gradually increased to reach a peak at day 3 of differentiation and decreased nearly to control levels at day 5 of differentiation (Figure 1G-H). In contrast, in shSH3KBP1-C2C12 cell, the transient increase of LC3B-II: ACTB was still detectable but markedly reduced and peaked earlier, at day 2 of differentiation. Nevertheless, we observed that both cell lines expressed MyH (myosin heavy chain), detectable as early as day 3 of differentiation (Figure S1E-F). The kinetics of MyH accumulation was statistically identical in both cell lines (figure S1F) confirming their efficient entry in the muscle differentiation program.

Taken together, our results indicate that SH3KBP1 is an actor of efficient basal autophagic activity and of the transient activation of autophagic flux occurring during muscle differentiation. However, when it is downregulated, the residual autophagic activity appears sufficient to support the early stages of myogenic differentiation.

### SH3KBP1 favors ER-phagy induction

Given that SH3KBP1 promotes basal autophagy in both myoblasts and myotubes and that ER-phagy contributes to ER-architecture remodeling in muscle cells [16], we next investigated whether SH3KBP1 is involved in the regulation of ER-phagy. This question was further supported by our previous observation that SH3KBP1 downregulation alters perinuclear ER organization in myotubes [23]. ER-phagy was first analyzed in shCTRL and shSH3KBP1 C2C12 myoblasts transfected with the fluorescent reporter ssRFP-GFP-KDEL [13] and treated or not with Rapa, a described pleiotropic inducer of ER-phagy [30]. ER-phagic activity was assessed by quantifying the number of RFP-positive vesicles per cell, which correspond to acidic lysosomal compartments containing ER-derived portions (Figure 2A). Under basal conditions (DMSO), shCTRL and shSH3KBP1 C2C12 cells displayed similar numbers of RFP-positive vesicles (Fig.2B), indicating that SH3KBP1 downregulation does not significantly affect basal ER-phagic activity. As expected, Rapa treatment induced a significant increase in the number of RFP-positive vesicles in shCTRL cells, consistent with an efficient ER-phagy induction (Figure 2A-B). In contrast, Rapa failed to significantly increase the number of RFP-positive vesicles in shSH3KBP1 cells compared with the corresponding DMSO condition, suggesting that SH3KBP1 downregulation impairs ER-phagy induction. A similar analysis was performed in 5 days-differentiated C2C12 myotubes, using loperamide (LOP), an ER-stress inducer known to activate ER-phagy [8,31] (Figure 2C-D). Consistent with our observations in myoblasts, basal ER-phagic activity was comparable between shCTRL and shSH3KBP1 myotubes under DMSO conditions. However, LOP treatment increased significantly the number of RFP-positive vesicles in shCTRL myotubes, whereas no significant increase was detected in shSH3KBP1 myotubes (Figure 2D). These results indicate that SH3KBP1 is required for an efficient ER-phagy induction in response to distinct stimuli, including rapamycin or loperamide, in both C2C12 myoblasts and myotubes.

**Figure 2.**
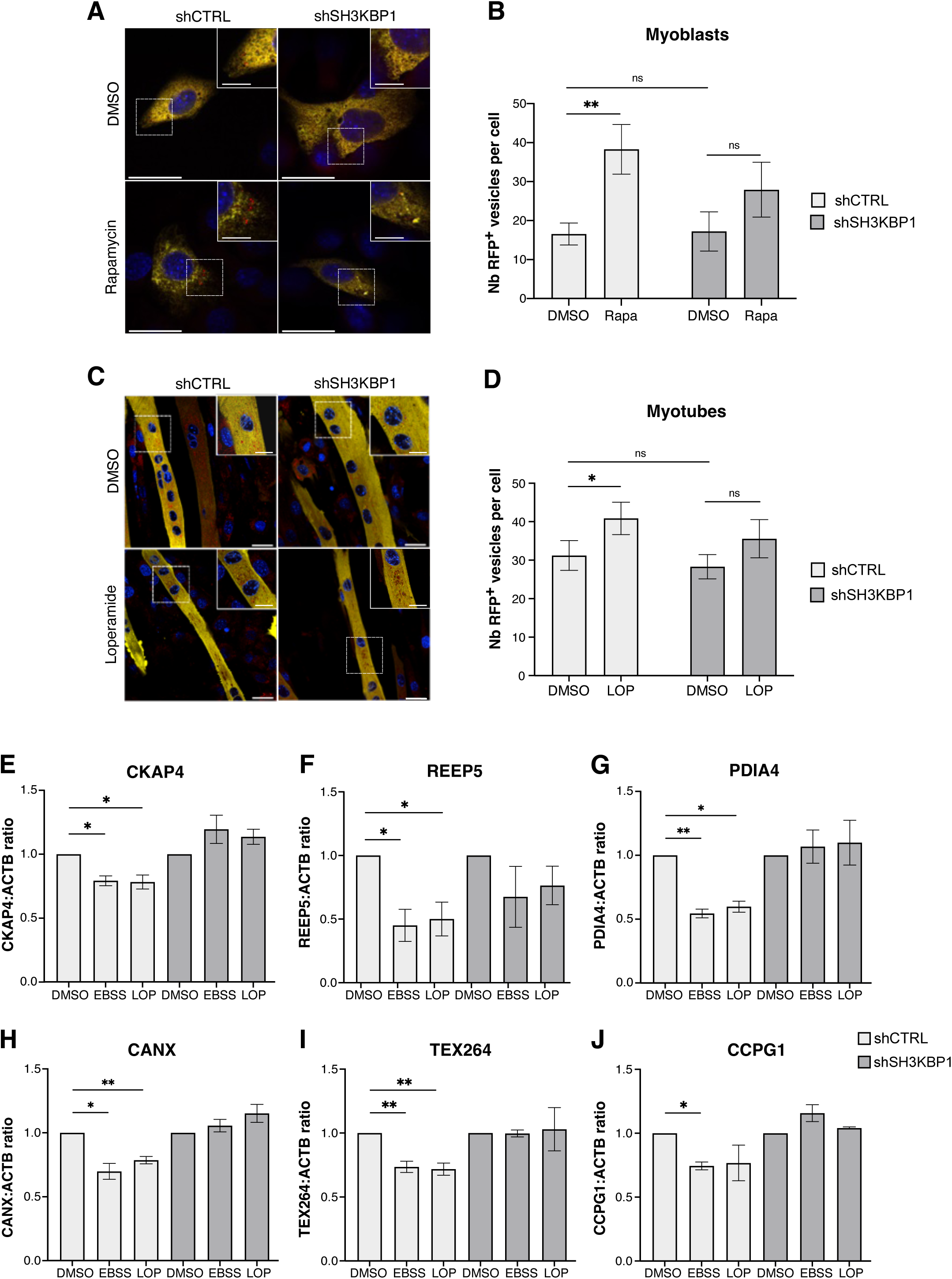
SH3KBP1 downregulation impairs ER-phagy induction. (**A and C**) Representative confocal images of C2C12 myoblasts (**A**) and 5 days-differentiated myotubes (**C**) stably expressing either control shCTRL or shSH3KBP1 and transiently transfected with the tandem fluorescent ER-phagy reporter ssRFP-GFP-KDEL. Myoblasts and myotubes were treated with DMSO (vehicle), Rapa (100 nM, 6 h) (**A**) or LOP (15 µM, 6h) (**C**). RFP-positive puncta correspond to ER-derived portions delivered to ALs. Scale bar: 30 µm (overviews images); 10μm (insets in A); 15μm (insets in C). (**B and D**) Quantification of ER-phagy activity by counting the number of RFP-positive puncta per cell, in shCTRL and shSH3KBP1 myoblasts (**B**) or myotubes (**D**). Data are presented as mean +/- SD; n=3 independent experiments. Statistical significance was assessed using two-way ANOVA. (**E to J**) Quantification of ER protein levels normalized to ACTB levels, following ER-phagy induction by EBSS or LOP treatment (15 μM) during 4 h, in shCTRL and shSH3KBP1 C2C12 cell lines: CKAP4 (**E**), REEP5 (**F**), PDIA4 (**G**), CANX (**H**), TEX264 (**I**) and CCPG1 (**J**). For corresponding western blots, see Figure S2A to S2F. Data are expressed as mean ± SD; n≥3 independent experiments. Statistical significance was assessed using Welch’s t-tests.

To support these observations, we quantified the levels of several ER-resident proteins in shCTRL and shSH3KBP1 myoblasts treated or not with LOP or subjected to starvation using Earle’s balanced salt solution (EBSS), two established ER-phagy inducers. In shCTRL cells, the ER-sheet structural protein CKAP4 was significantly reduced after both treatments compared to DMSO condition, in accordance with the degradation of ER portions upon ER-phagy induction. By contrast, CKAP4 levels remained unchanged in shSH3KBP1 cells (Figure 2E, Figure S2A). A similar observation was performed with the tubular ER protein REEP5 (Receptor expression-enhancing protein 5) whose levels decreased after EBSS or LOP treatment in shCTRL cells, whereas no significant declines were observed in shSH3KBP1 cells (Figure 2F, Figure S2B). These results indicate that SH3KBP1 downregulation impairs the degradation of both ER-sheets and ER-tubules compartments upon ER-phagy induction. The same analysis was performed with the ER chaperones PDIA4/ERp72 (protein disulfide isomerase family A member 4/ endoplasmic reticulum resident protein 72) and CANX, the latter being previously identified as an interactor of SH3KBP1 [23]. Both protein levels were significantly reduced following ER-phagy induction in shCTRL cells, but remained unchanged in shSH3KBP1 cells (Figure 2G-H, Figure S2C-D). Finally, we examined the turnover of ER-phagy receptors, such as TEX264 (testis expressed 264) and CCPG1 (cell cycle progression 1). Their levels were significantly reduced upon starvation and LOP treatment in shCTRL cells, whereas no significant changes were observed in shSH3KBP1 cells (Figure 2I-J, Figure S2E-F). Taken together, our results indicate that SH3KBP1 downregulation impairs the turnover of multiple ER-resident proteins involved in ER architecture, protein quality control and selective autophagy. These findings support a role for SH3KBP1 in ER-phagy induction by both pleiotropic or ER-stress inducers.

### SH3KBP1 shares the characteristics of soluble ER-phagy receptors

Given that SH3KBP1 modulates ER-phagy activity and the turnover of ER proteins, and that it has been reported to interact with some ER-proteins [19,23], we investigated whether SH3KBP1 could function as an ER-phagy receptor. We thus examined if SH3KBP1 displays key features of ER-phagy receptors, starting with the analysis of its localization to the ER. To this end, we performed co-immunostaining analyses of SH3KBP1 (red) and CKAP4 (green), described to localize mainly at ER sheets but also to a less extent at ER tubules [32] and previously reported to interact with SH3KBP1 [19]. Co-localization between SH3KBP1 and CKAP4 was observed in both C2C12 myoblasts and myotubes (Figure 3A-B), although the proteins displayed distinct distribution patterns. In myoblasts, CKAP4 showed a predominant perinuclear organization associated to a diffuse cytoplasmic signal, whereas SH3KBP1 exhibited a more aggregated distribution, concentrated at one side of the nucleus; accordingly, co-localization was mainly detected in the perinuclear region (Figure 3A). In myotubes, CKAP4 displayed both diffuse and punctate staining within the cytoplasm and around nuclei. SH3KBP1 was also detected as cytoplasmic and perinuclear puncta. Co-localization was therefore observed at the perinuclear region but also in elongated cytoplasmic structures (Figure 3B). These results thus suggest that SH3KBP1 associates with ER membranes, likely at their cytosolic face. Since SH3KBP1 has been reported to interact with CKAP4 in HeLa cells [19], we next investigated whether it associates with both CKAP4 and LC3B in C2C12 myoblasts, as expected for an ER-phagy receptor. We also examined whether these interactions are influenced by the modulation of autophagic activity. Co-immunoprecipitation experiments were performed using GFP-Trap magnetic particles, in C2C12 myoblasts expressing GFP-tagged full length SH3KBP1 (GFP-SH3KBP1_FL). The cells were either untreated (NT) or submitted to ER-phagy stimulation by starvation (EBSS) or LOP treatment, both combined with BafA1 treatment (EBSS+BafA1 and LOP+BafA1) to prevent degradation of the newly formed protein complexes (Figure 3C). Under basal conditions (DMSO), CKAP4, LC3B-I and LC3B-II were detected in the immunoprecipitated fraction, indicating that SH3KBP1 interacts with both CKAP4 and LC3B. These results confirm the previous reported interaction between SH3KBP1 and CKAP4 in HeLa cells and demonstrate, for the first time, an association between SH3KBP1 and LC3B in muscle cells. These interactions were also detected following EBSS+BafA1 or LOP+BafA1 treatments; interestingly, the amount of CKAP4 co-immunoprecipitated with GFP-SH3KBP1, normalized to the level of trapped GFP-SH3KBP1, was statistically increased after EBSS+BafA1 treatment and showed an increasing trend after LOP+ BafA1 treatment (Figure 3D). These results thus indicate that the association between SH3KBP1 and CKAP4 is enhanced upon autophagy and ER-phagy induction and suggest a regulated recruitment or stabilization of SH3KBP1 at the ER during this process. A similar pattern was observed for LC3B-II, the amount of LC3B-II detected in the immunoprecipitated fractions, normalized to trapped GFP-SH3KBP1 levels, was increased upon both treatments compared to NT condition (Figure 3C). This increase was statistically significant in EBSS/BafA1-treated cells; a non-significant upward trend was also observed in LOP+BafA1-treated cells (Figure 3E). The enhanced association between SH3KBP1 and LC3B-II after autophagy stimulation supports a functional link between SH3KBP1 and the autophagic machinery.

**Figure 3.**
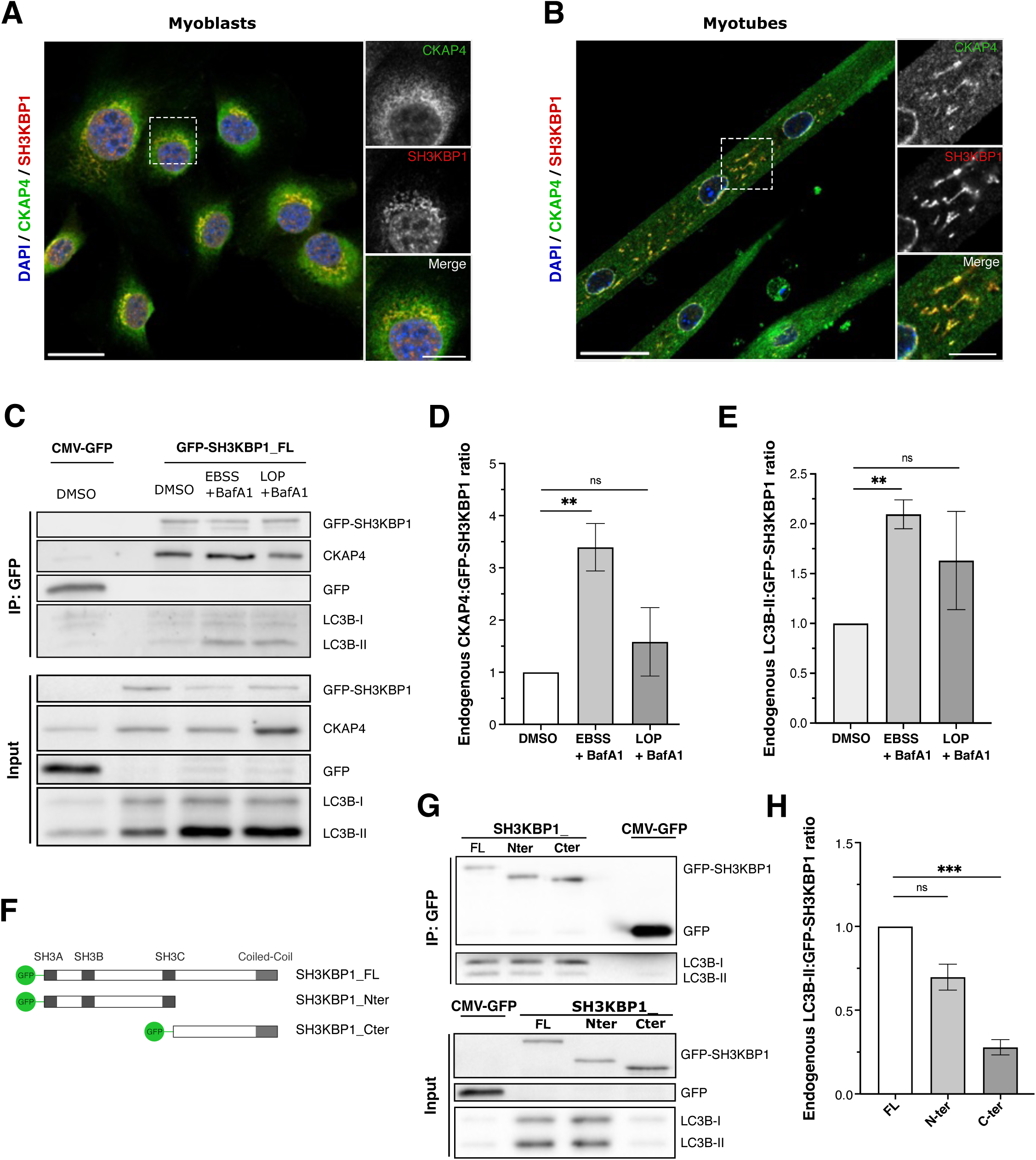
SH3KBP1 associates to ER and autophagosomes. (**A and B**) Representative confocal images of C2C12 myoblasts (**A**) and 5 days-differentiated myotubes (**B**) co-immunostained for SH3KBP1 (red) and CKAP4 (green). Nuclei were stained with DAPI (blue). Scale bar: 30 µm (overviews images); 10μm (insets in A and B). n=4 independent experiments (**C**) Representative western blot of co-immunoprecipitation experiments performed using GFP-Trap magnetic particles in C2C12 myoblasts transiently transfected with either CMV-GFP vector or the expression vector for full-length GFP-tagged-SH3KBP1 (SH3KBP1_FL). The day after, cells were either left untreated (DMSO) or submitted to autophagy induction by EBSS combined to BafA1 (100 nM) or by LOP (15 µM) combined to BafA1 for 4 hours. Input and immunoprecipitation (IP) fractions were analyzed with GFP, CKAP4 and LC3 antibodies. (**D**) Quantification of CKAP4-SH3KBP1 complexes assessed by measuring CKAP4 levels in the IP-fraction, normalized to immunoprecipitated GFP-SH3KBP1 (IP-fraction). Data are presented as mean +/- SD; n=5 independent experiments. (**E**) Quantification of LC3B-II-SH3KBP1 complexes as in (D); n=4 independent experiments. (**F**) Representative western blots of GFP-Trap experiments performed in C2C12 cells transfected with pCMV-GFP or expression vectors of GFP-tagged full-length SH3KBP1 (SH3KBP1_FL), GFP-tagged N-ter part of SH3KBP1 (SH3KBP1_Nter) or GFP-tagged C-ter part of SH3KBP1 (SH3KBP1_Cter). GFP and LC3B were revealed in the IP and input fractions. (**G**) Quantification of SH3KBP1-LC3B-II complexes as shown in (D). Data are presented as mean +/- SD; n=5 independent experiments. Statistical significances were assessed using Welch’s ANOVA tests, followed by Games-Howell post hoc tests.

To further delineate the molecular basis of this interaction, we performed in silico structural analyses using AlphaFold-based predictions of the interaction between SH3KBP1 and LC3B [33], and identified two putative regions within SH3KBP1 sequence. A first potential interaction site was predicted within the C-ter region of SH3KBP1 and a second in the N-ter region (Figure S3A). In addition, analyses, using iLIR database, revealed the presence of several putative LIR motifs throughout SH3KBP1 protein sequence (Figure S3B [34]. We thus examined which region of SH3KBP1 contributes to the binding to LC3B-II in C2C12 cells, using full length (SH3KBP1_FL), N-ter (SH3KBP1_Nter) or C-ter (SH3KBP1_Cter) halves of SH3KBP1 protein (Figure 3F) to perform GFP-Trap experiments (Figure 3 G and H). Both truncated forms were able to trap LC3B-II (Figure 3G) confirming Alpha-Fold and iLIR Database predictions that several LC3-binding sites may be present in SH3KBP1 protein. However, the amount of LC3B-II associated with either N-ter or C-ter construct was reduced compared to SH3KBP1_FL (Figure 3G), indicating that optimal association requires the full-length protein. This reduction reached statistical significance for the C-terminal construct, whereas the N-terminal construct showed a downward trend (Figure 3H), suggesting that both regions contribute to LC3-II binding, with a greater binding efficiency for the N-ter part. Altogether, these findings indicate that SH3KBP1 acts at the interface between the ER and the autophagic machinery; it associates with ER membranes via binding to CKAP4, while interacting with APs via binding to LC3B-II. This dual interaction supports a model in which SH3KBP1 function as a molecular adaptor linking ER-derived portions to the autophagic machinery during ER-phagy.

We next assessed whether SH3KBP1 could be targeted to autophagic compartments. Co-immunofluorescence analyses were performed using antibodies directed against SH3KBP1 (red) and the autophagosomal marker LC3B (green) in both C2C12 myoblasts and differentiated myotubes (Figure 4A and B, respectively). Under basal conditions, SH3KBP1 partially co-localized with LC3-positive puncta in both cell types. Quantitative analyses revealed that BafA1 treatment of myoblasts and myotubes significantly increased the number of SH3KBP1/LC3B colocalization events per cell, compared to control conditions. Moreover, autophagy induction by starvation, in the presence of BafA1 (EBSS+BafA1), further enhanced SH3KBP1 and LC3B co-localization, indicating that the association of SH3KBP1 to autophagosomal structures increases upon autophagy and ER-phagy stimulation. We then asked whether SH3KBP1 could also be detected in lysosomal compartments. Co-immunofluorescence analyses were performed using antibodies directed against SH3KBP1 and the lysosomal marker LAMP1 (lysosome associated membrane protein 1), in C2C12 myoblasts and myotubes (Figure 4C and D, respectively). Basal colocalization between SH3KBP1 and LAMP1 was observed in both cell types under control conditions. Quantifications revealed that ER-phagy induction by EBSS or LOP significantly increased SH3KBP1-LAMP1 co-localization in both myoblasts and myotubes. These observations indicate that SH3KBP1 is present within lysosomal compartments and that its association with these structures increases upon autophagy induction. Together, the increased association of SH3KBP1 with both autophagosomal and lysosomal compartments upon autophagy modulation suggests that SH3KBP1 may itself be degraded by autophagy. To further confirm this point, SH3KBP1 levels were analyzed by western blot in C2C12 myotubes, upon autophagy/ER-phagy modulation. As expected, LOP treatment induced a progressive increase of LC3B-II (Figure 4E), confirming activation of the autophagy/ER-phagy process. In contrast, LOP treatment led to a time-dependent decrease of SH3KBP1 level (Figure 4E), which became statistically significant after 6h of treatment (Figure 4F). Conversely, BafA1 treatment resulted in a progressive accumulation of LC3B-II, consistent with the inhibition of autophagic degradation and was associated to accumulation of SH3KBP1 (Figure 4G), with a significant increase detected as early as 3 hours of treatment (Figure 4H). Hence, our observations demonstrate that SH3KBP1 is recruited to LC3B-II-positive APs and addressed to LAMP1-positive lysosomal/autolysosomal compartments to be degraded, confirming that SH3KBP1 protein turnover is controlled by autophagy.

**Figure 4.**
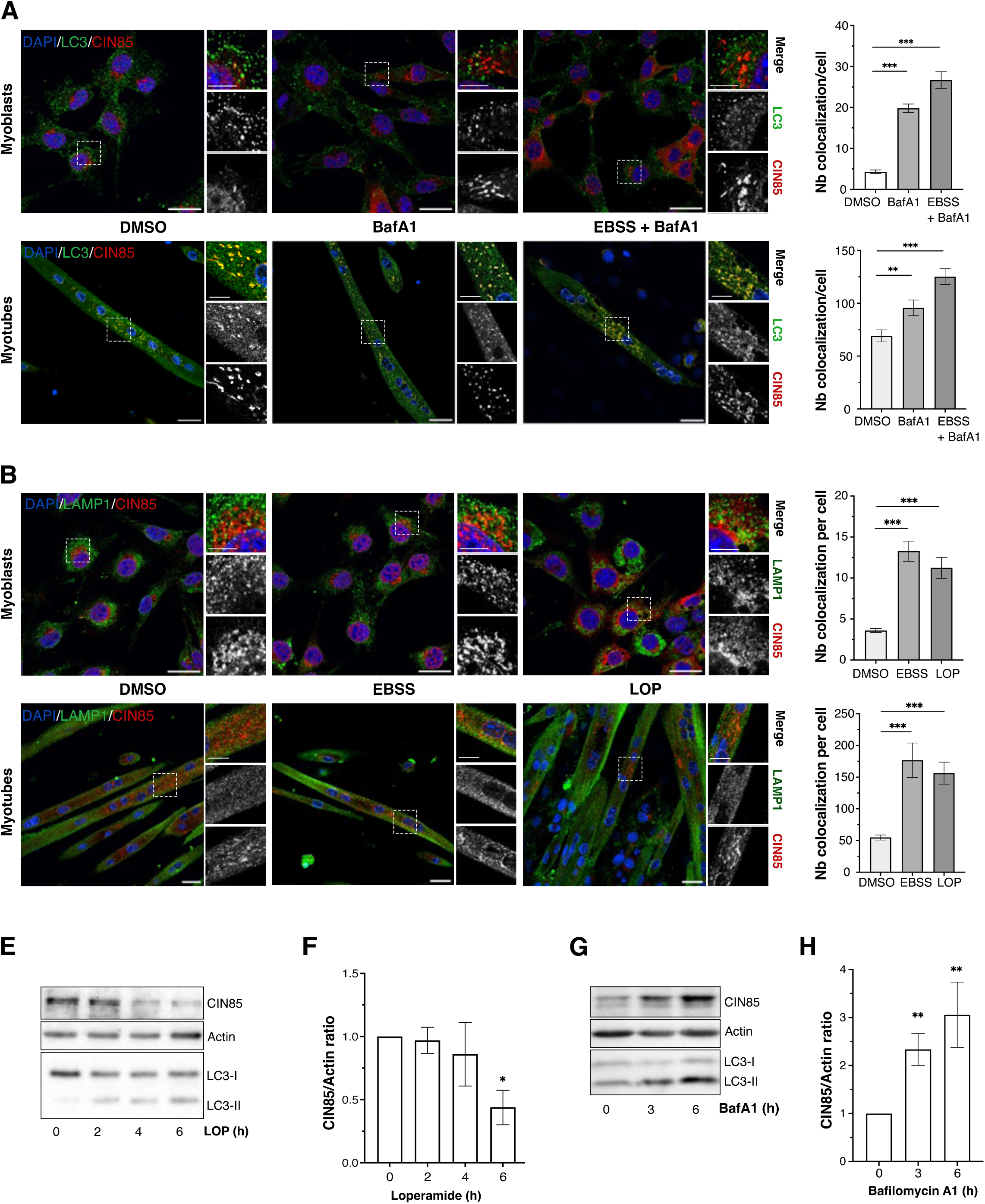
SH3KBP1 is degraded by ER-phagy. (**A, B, C and D**) Representative confocal images of SH3KBP1 (red) co-immunostained with LC3B (green, **A**) or LAMP1 (green, **B**) in C2C12 myoblasts (**A** and **C**) and 5 days-differentiated myotubes (**B** and **D**). Cells were left untreated (DMSO, vehicle) or treated with BafA1 (100 nM, 6 h) or EBSS+BafA1 (6 h) (**A** and **B**) or with EBSS (4h) or LOP (15 µM, 6 h) (**C** and **D**). Nuclei were stained with DAPI (blue). Scale bar: 30 µm (overviews images); 10μm (insets). Quantifications show the number of SH3KBP1-LC3B (**A** and **B**) or SH3KBP1-LAMP1 (**C** and **D**) colocalization events per cell. Data are presented as mean +/- SD; n=3. Statistical significance was assessed using Kruskal-Wallis test followed by Dunn’s multiple comparisons test. (**E and F**) Representative western blot showing SH3KBP1, LC3B and ACTB (loading control) in C2C12 cells treated with LOP (15 µM, 0 to 6 h) (**E**), and the corresponding quantification of SH3KBP1 levels normalized to ACTB levels (**F**). Data are presented as mean +/- SD; n=3 independent experiments. Statistical analysis was performed using Welch’s ANOVA followed by Games-Howell multiple comparisons test. (**G and H**) Representative western blot showing SH3KBP1, LC3B and ACTB (loading control) in C2C12 cells treated with BafA1 (100nM, 0 to 6h) (**G**), and the corresponding quantification of SH3KBP1 levels normalized to ACTB levels (**H**). Data are presented as mean +/- SD; n=5 independent experiments. Statistical analysis was performed using Kruskal-Wallis followed by Dunn’s multiple comparisons test.

Altogether our results demonstrate that SH3KBP1 binds to LC3B-II via its LIR motifs and associates to the ER through its interaction with CKAP4; moreover, it is itself degraded upon ER-phagy stimulation. We thus propose that SH3KBP1 is a novel soluble ER-phagy receptor that regulate ER-phagy activity in muscle cells. Given its ubiquitous expression and involvement in membrane trafficking pathways, SH3KBP1 may also contribute to ER-phagy regulation in other cellular contexts.

## Discussion

ER-phagy selectively targets portions of the ER, for degradation by lysosomes. This process participates to the regulation of ER shape, size and function and relies on the activation of ER-phagy receptors. Several ER-phagy receptors have now been described [8]. The majority are transmembrane ER-resident proteins including members of the RETREG/FAM134 family [35], as well as CCPG1 [36], TEX264 [37], SEC62 (SEC62 preprotein translocator factor) [38], RTN3L (reticulon 3 long form) [39] and ATL3 (atlastin GTPase 3) [40]. Although less numerous, soluble ER-phagy receptors have also been discovered (CALCOCO1 [calcium-binding and coiled-coil domain-containing protein1], CDK5RAP3/C53 [CDK5 regulatory subunit associated protein 3], EPR1 (effector cell peptidase receptor 1) or SQSTM1/p62 [sequestosome 1]) [12,41–43]. These receptors rely on interactions with ER-resident proteins such as VAP (Ventilator-associated pneumonia) proteins for CALCOCO1 and Epr1, UFL1 (UFM1 specific ligase 1) for CDK5RAP3/C53 or ubiquitin for SQSTM1/p62. In this study, we have identified SH3KBP1 as a novel soluble ER-phagy receptor required for efficient ER-phagy induction by pharmacological treatment, starvation and ER-stress, in both myoblast and myotubes. Unlike most of the other ER-phagy receptors whose under-expression does not significantly affect basal autophagy flux, SH3KBP1 downregulation reduced AP formation and maturation. These results are consistent with our previous work showing reduced LC3B-II accumulation upon BafA1 treatment in shSH3KBP1 C2C12 cells, as well as increased LC3BII-positives myofibers, in mice *tibialis anterior* (TA) muscles, following in vivo shRNA-mediated SH3KBP1 knockdown [23]. In contrast, rapa-induced bulk autophagy was not altered when SH3KBP1 was downregulated, suggesting that SH3KBP1 primarily contributes to basal autophagic machinery rather than maximal inducible autophagy. Similar observations were also reported for the soluble ER-phagy receptor CALCOCO1 [12] raising the possibility that this differential regulation of basal and bulk autophagy may be a feature of soluble ER-phagy receptors, although further investigations are required.

We then demonstrated that SH3KBP1 downregulation impairs ER-phagy induction by Rapa, LOP or starvation with EBSS. Rapa inhibits MTOR kinase, whereas EBSS induces amino-acid starvation and glucose restriction; both act as pleiotropic ER-phagy inducers, as they also activate bulk autophagy and other forms of organellophagy [8]. LOP induces ER stress through ATF4 (activating transcription factor 4) activation [31], and is also described to modulate calcium channels, being more an ER-centric inducer [44]. Our results thus suggest that SH3KBP1 functions as a broad stress-responsive ER-phagy receptor activated by diverse stimuli. We also confirmed the involvement of SH3KBP1 in the induction of ER-phagy by observing that SH3KBP1 regulates the turnover of various ER-resident proteins such as CKAP4, REEP5, PDIA4, CANX, TEX264 and CCPG1, as already observed for other soluble ER-phagy receptors [12,41]. These proteins include transmembrane (CANX) or luminal (PDIA4) chaperones, ER sheets associated markers (CKAP4) or tubular ones (REEP5, TEX264, CCPG1), architectural proteins (CKAP4, REEP5) or ER-phagy receptors themselves (TEX264, CCPG1). Our results thus suggest that SH3KBP1-mediated ER-phagy is not restricted to specific ER region but rather affect global ER remodeling.

Given that SH3KBP1 is distributed across several membrane trafficking compartments including perinuclear region, Golgi, ERGIC and endosomes [21] and being able to stimulate ER-phagy activity, we examined its subcellular localization relative to the transmembrane ER protein CKAP4, predominantly associated with ER sheets but also detected in peripheral ER tubules [45]. SH3KBP1 and CKAP4 co-localized predominantly in the perinuclear region but also in cytoplasmic patches in myoblasts and in elongated structures in myotubes. These cytoplasmic puncta may correspond to endosomal compartments, consistent with previous studies showing occasional CKAP4 trafficking from the ER to endosomes [46,47]. Alternatively, they may reflect liquid-liquid phase separation (LLPS) of SH3KBP1 and CKAP4 [48], as SH3KBP1 possesses 4 intrinsically disordered region (IDRs), a type of regions known to efficiently promote LLPS [49]. This latter hypothesis is further supported by a previous report describing SH3KBP1 engagement in phase separated-condensates with SLP65 protein (Src homology2 domain-containing leucocyte protein of 65 kDa), during B cell receptor signaling [50]. The elongated structures observed in myotubes may reflect SH3KBP1 colocalization with CKAP4 to the nascent sarcoplasmic reticulum (SR) during differentiation. Our imaging and co-immunoprecipitation data thus support the existence of a SH3KBP1-CKAP4 complex in muscle and are consistent with previous reports performed in other cell types [19].

Of interest, GFP-SH3KBP1 immunoprecipitation not only recovered CKAP4 but also LC3 cytosolic form LC3B-I and its lipidated form LC3B-II. Moreover, SH3KBP1 interactions with both CKAP4 and LC3B-II were increased upon autophagy/ER-phagy stimulation, demonstrating a dynamic recruitment of SH3KBP1 to both ER and APs during ER-phagy and reinforcing our proposal that SH3KBP1 is a soluble ER-phagy receptor. Truncation analyses revealed that both the N-ter and C-ter regions of SH3KBP1 contribute to LC3B binding, with a stronger contribution from the N-ter region, in agreement with AlphaFold-based predictions [33]. In contrast, deletions of high-confidence LIR motifs predicted in the N-ter region by the iLIR database (PSSM-based selection; data not shown) did not significantly affect LC3B-II binding, suggesting that LC3B-II interaction relies on a multivalent interface rather than a single dominant canonical LIR motif. This may reflect a compensatory and/or cooperative contribution of the six canonical LIR motifs (W/F/YxxL/V/I) identified in the N-ter part of SH3KBP1 together with the extended LIR motif localized in its C-ter part (iLIR database [34]). Indeed, interactions between autophagy receptors and ATG8 protein family (GABARAP and LC3 families) often rely on multivalent binding mechanisms involving multiple LIR motifs rather than a single dominant site [51].

SH3KBP1 protein sequence also contains short IDR in its N-ter region and long IDR in its C-ter part. IDRs, like presence of LIR motifs, are increasingly recognized as key elements in membrane-bound ER-phagy receptors promoting phase separation, ER-phagy receptors clustering, ER-membrane deformation [49] and fragmentation [14]. Their presence further supports the classification of SH3KBP1 as a soluble ER-phagy receptors, especially since IDR regions are also reported in the other soluble ER-phagy receptors identified so far (CALCOCO1, CDK5RAP3, EPR1, SQSTM1). Whether IDR of soluble ER-phagy receptors have similar functions to those present in membrane-bound ER-phagy receptors remains speculative but represents an interesting point to future investigations.

Finally, we demonstrated that SH3KBP1 is targeted to autophagosomes and lysosomes and undergoes autophagic degradation. SH3KBP1 levels decreased upon LOP treatment, whereas BafA1 prevented this degradation and led to protein accumulation. In contrast, no accumulation of SH3KBP1 was observed upon treatment with MG132, a proteasome inhibitor (data not shown), further supporting an autophagy-dependent turnover of SH3KBP1. Together, these data establish SH3KBP1 as a soluble ER-phagy receptor. The pending question is how ER-phagy receptors and in particular SH3KBP1 are activated to induce ER-phagy. Indeed, SH3KBP1 is a scaffold protein that promotes the formation of large macromolecular assemblies and can participate in different signaling pathways. As for other scaffold proteins, post-translational modifications and/or interaction with specific effectors may contribute to its regulation. Of interest, autophagy receptors such as the RETREG family are regulated through phosphorylation and ubiquitination processes [13]. Regarding SH3KBP1, phosphorylation and mono-ubiquitination have already been described, notably in response to EGFR (epidermal growth factor receptor) signaling [52] [53] and multiple phosphorylation sites have been identified. More recently, phosphorylation of SH3KBP1 at residue S230 was also shown to regulate its ability to undergo liquid-liquid phase separation with SLP65, by controlling SH3KBP1 intramolecular interaction between SH3 and proline rich domains [50]. It is therefore tempting to postulate that similar post-translational modifications could regulate SH3KBP1 activity in response to ER-phagy inducers. Another open question is whether SH3KBP1-dependant ER-phagy contribute to ER architecture but also to degradation of specific ER-luminal proteins, either through interaction with CKAP4 alone or through a tripartite complex involving CKAP4 and CANX. Indeed, we previously demonstrated that SH3KBP1 interacts CANX via its C-ter region [23]. CANX is an ER-chaperone that, together with its partner CALR (calreticulin), senses misfolded proteins and has been shown to participate in the ER-phagy mediated degradation of pro-collagen via ER-phagy [54].

Regarding the contexts in which SH3KBP1-dependent ER-phagy operates, we detected this pathway in both myoblasts and myotubes. More specifically, we detected an increased accumulation of APs during the differentiation process, consistent with reports of autophagy activation during myogenesis [16]. However, this increase was attenuated in SH3KBP1-underexpressing myoblasts, indicating that SH3KBP1-dependent autophagy contributes to myogenic differentiation. In a previous study, we demonstrated that commitment into muscle differentiation is not affected by SH3KBP1 downregulation; however, an increase in fusion activity was observed. Nonetheless, SH3KBP1-underexpressing myotubes were able to accumulate the myogenic marker MyH, indicating an efficient progression through differentiation; however, myotubes displayed a disorganized perinuclear ER [23] and, at myofiber stage, altered SR/triad organization, hence highlighting the importance of SH3KBP1 and ER-phagy in maintaining ER/SR architecture [23]. The role of ER-phagy in the regulation of ER dynamics during myogenesis has also been nicely demonstrated by Buonomo and colleagues who showed that RETREG1 (reticulophagy regulator 1) family and notably the B2 isoform is a key ER-phagy receptor during ER to SR remodeling [16] [55]. SH3KBP1 might maximize the ER-phagy process governed by RETREG1, since we observed an attenuated ER-phagy induction when SH3KBP1 is downregulated. ER-phagy has also been shown to be crucial during neurogenesis, where ER architecture needs to generate highly refined tubular network in dendrites and axons. Whether SH3KBP1 could play a similar role in this context would be interesting to find out. Finally, we previously demonstrated that under-expression of SH3KBP1 in the TA muscle of a wild type mice induces an atrophic phenotype. Interestingly, when performed in the TA muscle of a mouse model of centronuclear myopathy (KI-Dnm2^R465W^), the atrophic phenotype was drastically exacerbated and associated with altered autophagic processes [23]. These observations thus suggest that SH3KBP1-dependant ER-phagy might be involved in the pathophysiology of centronuclear myopathies. Understanding the role of SH3KBP1 potential as an ER-shaping factor as well as its modes of activation in different tissue contexts may provide valuable insight for the development of therapeutic strategies targeting diseases associated to a deficient ER-phagy, including neurological and muscle disorders, cancer and infections [8].

## Materials and Methods

### Plasmids and reagents

pCMV-GFP was a gift from Connie Cepko (Addgene, plasmid # 11153; [56]). The mRFP-GFP-LC3 plasmid (ptfLC3) was a gift from Tamotsu Yoshimori (Addgene plasmid # 21074; [24]) and the ssRFP-GFP-KDEL plasmid was kindly gifted by A. Stolz (Goethe University, Frankfurt am Main, Germany; [13]). GFP-SH3KBP1 constructs were previously described (provided by Dr V. Buchman; [18]). Briefly, GFP was added at the N-ter of the following constructs: full-length (FL) SH3KBP1 (SH3KBP1_FL, amino acids 1 to 665), N-ter fragment (SH3KBP1_Nter, amino acids 1-354) and C-ter fragment (SH3KBP1_Cter, amino acids 328-665).

The following reagents were used to induce autophagy and/or ER-phagy: Earle’s Balanced Salt Solution (EBSS, [Gibco, 24010043]), Rapamycin (UBPBio, F6130; 100 nM), Loperamide hydrochloride (Enzo Life Science, ALX550253G005; 15 µM,). BafA1 (MedChemExpress, HY-100558; 100 nM,) was used to block the autophagic flux.

### Cell lines, cell culture and transfections

C2C12 is a commonly used myoblast cell line [57] [58]. C2C12 cell lines expressing control shRNAs (shCTRL, Mission pLKO.1-Puro non mammalian shRNA control plasmid, [Merck, SHC002]) or shRNAs directed against SH3KBP1 (shSH3KBP1, Mission® shRNA, [Merck, clone TRCN0000088508]) have already been described [23].

Myoblasts were cultured in Dulbecco’s Modified Eagle Medium (DMEM) supplemented with 15% fetal bovine serum (FBS) (Gibco, 26140079) and 1% penicillin-streptomycin (Gibco, 15140122; 100 U/mL) in humidified incubator at 37°C and under 5% of CO_2_. To induce myogenic differentiation, cells were seeded at 80% confluence, and, the day after, switched to DMEM differentiation medium containing 2% horse serum (Gibco #16050122) and 1% penicillin-streptomycin (100U/mL), as described by McMahon et al. [58].

Transient transfections were performed using JetOptimus (Sartorius jetOPTIMUS®, 101000006) reagents according to manufacturer’s instructions.

### Co-immunostainings

Cells were seeded in µ-slide 4-wells (1.5 to 2.5×10^4^ cells/well) or 8-wells (0.75×10^4^ cells/well) chambers (Ibidi, 80806 or 80426, respectively). For myotube experiments, cells were switched to differentiation medium 24 h after seeding and maintained for 5 days as described above. Following the indicated treatments, cells were rinsed with warm phosphate buffered saline (PBS, [Euromedex, ET330]), fixed in 4% paraformaldehyde (PFA) for 20 min at 37°C and permeabilized with 0.01% Triton X-100 in PBS for 10 min at room temperature. For indirect immunofluorescence, cells were incubated overnight at 4°C with primary antibodies directed against SH3KBP1 (Atlas, HPA#003355; 1:250), either alone or in combination with LC3 (NanoTools, 0231-100/LC3-5F10; 1/200) or LAMP1 (Proteintech, 67300-1-Ig; 1/500). After washing, cells were incubated during 1 h in secondary antibody (Invitrogen; A21236, A11029; 1:5000) in the presence of 4’,6-diamidino-2-phenylindole (DAPI) (Life Technologies, D1306; 1X, 50µg/mL). For experiments using conjugated-antibodies, SH3KBP1 immunostaining was followed by incubation with Coralite488-conjugated antibodies against CKAP4/CLIMP63 (Proteintech, CL488-16686; 1/500), LC3 (Proteintech, CL488-14600; 1/500) or LAMP1 (Proteintech, CL488-67300; 1/500). After several washings, cells were stored at 4°C in PBS containing 15 µM sodium azide, until image acquisition. Because of differences in antibody accessibility and epitope detection between myoblasts and myotubes, staining strategies were optimized for each antibody combination and cell type.

### Confocal microscopy and image analysis

Confocal imaging was performed on a Nikon AX R microscope (Eclipse Ti2-E platform) equipped with a 60x oil-immersion objective (CFI Plan Apochromat Lambda, NA 1.4, Nikon). Single Z-slice images were acquired with the confocal pinhole set to 1 Airy unit, yielding a submicron optical section. For each condition, 10 randomly selected fields per well were recorded.

All images were acquired using manufacturer-supplied software (Nikon) and processed with Fiji (NIH). Background subtraction and mild filtering were applied prior to threshold-based segmentation. Colocalization was quantified using binary mask-based image multiplication (logical AND) between SH3KBP1 and LC3 or LAMP1 signals. For myoblasts, quantifications were performed at the single-cell level, using DAPI staining to define nuclei as individual cellular units. For myotubes, quantifications were performed on individual multinucleated myotubes, with fields of view selected during image acquisition to include a single centrally positioned myotube whenever possible.

### Immunoprecipitation experiments (GFP-Trap)

Immunoprecipitation experiments were performed using ChromoTek GFP-Trap® magnetic particles (Proteintech, gtd-20). Cells were seeded at 3.5×10^5^ to 5×10^5^ cells per 100 mm culture dishes and transiently transfected with either control vector or GFP-SH3KBP1 constructs (FL, N-ter or C-ter) using JetOptimus reagent (as described above). 24h post-transfection, cells were lysed in ice-cold lysis buffer (10 mM Tris HCl [Sigma-Aldrich, T1503] pH 7.5, 150 mM NaCl [Euromedex, 1112-A], 0.5 mM EDTA [Sigma-Aldrich, EDS] pH8, 0.5% Nonidet P40 substitute [ThermoFisher Scientific, J19628.K2]) supplemented with Complete™ protease inhibitor cocktail (Roche, 11836170001; 1X). Lysates were clarified by centrifugation (17,000 g, 10 min, 4°C), and supernatant were incubated with GFP-Trap beads for 2h at 4°C under rotation. An aliquot of the lysate was kept as input. Beads were washed three times with dilution buffer (10 mM Tris HCl pH 7.5, 150 mM NaCl, 0.5 mM EDTA pH8, 1X protease inhibitor), and bound proteins were eluted in Laemmli buffer, boiled at 95°C for 10 min, and analyzed by SDS PAGE and immunoblotting.

### Protein sample preparation and Western BloĖng

Protein extracts were prepared in Radioimmunoprecipitation assay buffer (RIPA) supplemented with protease and phosphatase inhibitors (Roche, 11836170001 and PHOSS-RO, respectively; 1X). Lysates were sonicated, clarified by centrifugation and protein concentrations were determined using a BCA Protein Assay (Thermo Scientific, 23225). Equal amounts of protein (15-30 µg) were separated by SDS-PAGE (10-15% gels or 4-15% gradient gels [BioRad, 4561083]) and transferred to nitrocellulose membranes (Amersham, GE10600001). Membranes were blocked and incubated with the following primary antibodies: anti-SH3KBP1 (Atlas, HPA003355; 1:10,000), anti-LC3B (Sigma-Aldrich, L7543; 1:1,000), anti-ACTB (Millipore, Mab1501; clone C4; 1:5,000) and anti-GFP (Chromotek, H39-100; 1:1,000), anti-MYH1E (DSHB, MF20; 1/500), anti-CKAP4 (Proteintech, 16686-1-AP; 1:1,000), anti-REEP5 (Proteintech, 14643-1-AP; 1:5,000), anti-PDIA4 (Proteintech, 66365-1-AP; 1:2,000), anti-CANX (Proteintech, 660903-1-Ig; 1:2,000), anti-TEX264 (Proteintech, 25858-1-Ig; 1:1000) and anti-CCPG1(Proteintech, 13861-1-AP; 1:500). After washing, HRP-(horseradish peroxidase) conjugated secondary antibodies (Amersham, NA9310V and NA9340V; 1:5,000) were applied and signal was detected using enhanced chemiluminescence (ECL Prime, [Amersham, GERPN2236]) and acquired with a Chemidoc imaging system (BioRad Chemidoc™ Imager). Quantifications were performed by Image Lab Software v6.0, which displays saturated pixels in a different color, in order that every quantification is performed with protein signals in the linear range of detection of the imaging system.

### Statistics

Graphs were generated using GraphPad Prism software. Statistical analyses were conducted in RStudio using the RCommander package. Normality was assessed using the Shapiro-Wilk test, and homogeneity of variances was tested with Bartlett’s or Brown-Forsythe’s tests as appropriate. For two-group comparisons, parametric (Student’s or Welch’s t-test) or non-parametric (Mann-Whitney or Wilcoxon rank-sum test) tests were used depending on data distribution. For comparisons involving more than two groups, parametric data were analyzed using one-way ANOVA followed by Tukey’s or Games-Howell post hoc tests, while non parametric data were analyzed using Kruskal-Wallis tests followed by Dunn’s multiple comparison tests. P-values < 0.05 were considered were considered statistically significant (* p<0.05, ** p<0.01, *** p<0.001).

## Acknowledgements

We thank the members of Gache and Strappazzon teams and Dr Vincent Jacquemond for helpful scientific discussions. We are grateful to Alexandra Stolz for the kind gift of ssRFP-GFP-KDEL plasmid and Marion Carpentier and Dr Flavie Strappazzon for their help with LC3 antibodies. We thank Dr Caroline Brun for her efficient help with Nikon AX R microscope. We thank Sidy Fall for his advice on statistical analyses.

## Disclosure statement

No potential conflict of interest was reported by the authors.

## Funding

This work was supported by the French National Agency (ANR-21-CE14-0064, CKR), the Association Française contre les Myopathies (AFM-Téléthon: Alliance MyoNeurALP2 program CKR and VG, project #2.2.2).

## Data availability statement

Original data used to perform quantifications shown in this study are available upon reasonable request to the corresponding author.

## Author contributions

CRediT. **Marine Daura**: Conceptualization, Formal analysis, Investigation, Methodology, Validation, Visualization, Writing-original draft; **Elina Vergara**: Investigation; **Leslie Andromaque**: Formal analysis, Investigation; **Agathe Leddet**: Investigation; **Emilie Christin**: Investigation; **Céline Malleval**: Formal analysis; **Vincent Gache**: Conceptualization, Funding acquisition, Resources, Writing-review & editing; **Carole Kretz-Remy**: Conceptualization, Funding acquisition, Investigation, Methodology, Project administration, Resources, Supervision, Validation, Visualization, Writing-original draft, Writing-review & editing.

## Abbreviations

ATG: Autophagy related
AL: autolysosome
AP: autophagosome
BafA1: bafilomycin A1
C-ter: C-terminal
DAPI: 4’,6-diamidino-2-phenylindole
DMSO: dimethyl sulfoxide
EBSS: Earle’s Balanced Salt Solution
ER: endoplasmic reticulum
FL: Full Length
GFP: green fluorescent protein
IDR: intrinsically disordered region
LAMP1: lysosomal-associated membrane protein 1
MAP1LC3/LC3: microtubule-associated protein 1 light chain-3
LIR: LC3 Interacting Region
LOP: loperamide hydrochloride
mTOR: mechanistic target of rapamycin kinase
mRFP: mammalian red fluorescent protein
MyH: Myosin heavy chain
N-ter: N-terminal
PBS: phosphate buffered saline
Rapa: rapamycin
shCTRL: control short hairpin RNA
shSH3KBP1: short hairpin RNA directed against SH3KBP1 mRNA
SR: sarcoplasmic reticulum
TA: tibialis anterior

## Notes

### Competing Interest Statement

The authors have declared no competing interest.

