## supplementary figures for "SH3KBP1/CIN85, a new actor of ER-phagy in muscle"

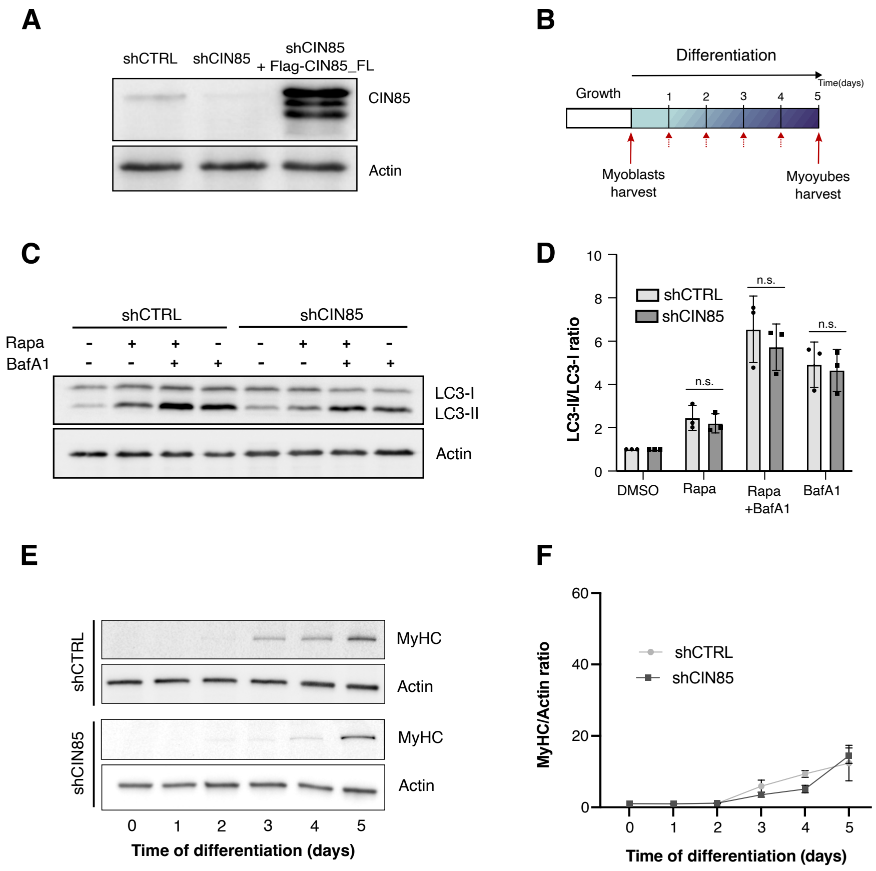


**Figure S1**. Validation of SH3KBP1 levels modulation and assessment of autophagy during C2C12 differentiation. (**A**) Representative western blot showing SH3KBP1 expression in shCTRL and shSH3KBP1 C2C12 myoblasts, as well as in shSH3KBP1 cells re-expressing full-length Flag-SH3KBP1 (Flag-SH3KBP1_FL). ACTB was used as a loading control. (**B**) Schematic representation of the C2C12 differentiation protocol. Cells were maintained in growth medium before switching to differentiation medium. Solid red arrows indicate endpoint harvests of myoblasts and myotubes, whereas dashed red arrows indicate daily sampling for kinetic analyses. (**C and D**) Representative western blot analysis of LC3B-II levels in shCTRL and shSH3KBP1 C2C12 myoblasts treated with rapamycin (Rapa, 100 nM), bafilomycin A1 (BafA1, 100 nM), or a combination of both treatment for 4h (**C**) and corresponding quantification calculated as the LC3B-II:LC3B-I ratio and normalized to the DMSO condition, which was set to 1 within each cell line (**D**). Data are presented as mean +/- SD; n=3 independent experiments. (**E and F**) Representative western blot showing myosin heavy chain (MyH) expression during differentiation of shCTRL and shSH3KBP1 C2C12 cells (**E**) and corresponding quantification normalized to ACTB (**F**). Data are presented as mean +/- SD; n=3 independent experiments.

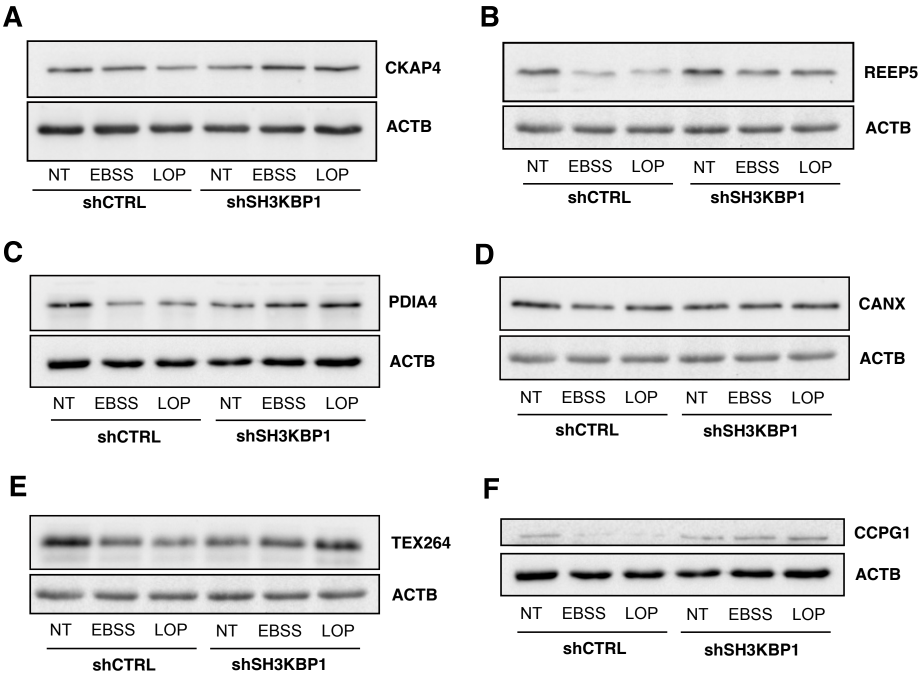


**Figure S2**: SH3KBP1 downregulation impairs the turnover of ER-resident proteins upon ER-phagy induction. (**A - F**) Representative western blots showing CKAP4 (**A**), REEP5 (**B**), PDIA4 (**C**), CANX (**D**), TEX264 (**E**) and CCPG1 (**F**) in shCTRL and shSH3KBP1 cells treated with DMSO, EBSS or loperamide (LOP, 15 μM) for 4h prior to protein extraction. ACTB was used as a loading control. Corresponding blot quantifications are shown in Fig. 2E-J.

**A**

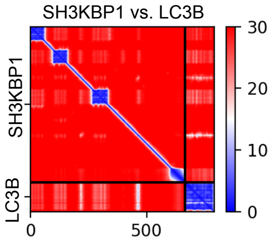


**B**

| Motif | Position (residues) | Pattern | PSSM Score |
|---|---|---|---|
| WxxL | 213-218 | VGFGDI | 4 |
| WxxL | 235-240 | NDFLPV | 12 |
| WxxL | 269-274 | KDYCKV | 10 |
| WxxL | 134-139 | GWWEGV | 10 |
| WxxL | 320-325 | DNFVKL | 13 |
| xLIR | 464-469 | EGFDSV | 11 |

**Figure S3**: In silico prediction of SH3KBP1 interaction motifs. (**A**) AlphaFold-based predicted aligned error (PAE) map showing the predicted interaction between SH3KBP1 and LC3B. (**B**) Summary of putative LC3-interacting region (LIR) motifs identified in SH3KBP1 protein sequence using the iLIR database tool; type, position, sequence, and PSSM (Position Specific Scoring Matrix) are indicated for each motif.
